# *Slc26a9^P2ACre^*, a new CRE driver to regulate gene expression in the otic placode lineage and other FGFR2b-dependent epithelia

**DOI:** 10.1101/2020.05.26.118026

**Authors:** Lisa D. Urness, Xiaofen Wang, Chaoying Li, Rolen M. Quadros, Donald W. Harms, Channabasavaiah B. Gurumurthy, Suzanne L. Mansour

**Author notes:** Corresponding Author Suzanne L. Mansour, Department of Human Genetics, University of Utah, 15 North, 2030 East, Room 5100, Salt Lake City, UT 84112-5330.

## Abstract

Pan-otic CRE drivers enable gene regulation throughout the otic placode lineage, comprising the inner ear epithelium and neurons. However, intersection of extra-otic gene-of-interest expression with the CRE lineage can compromise viability and impede auditory analyses. Furthermore, extant pan-otic CREs recombine in auditory and vestibular brain nuclei, making it difficult to ascribe resulting phenotypes solely to the inner ear. We previously identified *Slc26a9* as an otic placode-specific target of FGFR2b ligands, FGF3 and FGF10. We show here that *Slc26a9* is otic-specific through E10.5, but not required for hearing. We targeted *P2ACre* to the *Slc26a9* stop codon, generating *Slc26a9*^*P2ACre*^ mice, and observed CRE activity throughout the otic epithelium and neurons, with little activity evident in the brain. Notably, recombination was detected in many FGFR2b ligand-dependent epithelia. We generated *Fgf10* and *Fgf8* conditional mutants, and activated an FGFR2b ligand trap from E17.5-P3. In contrast to analogous mice generated with other pan-otic CREs, these were viable. Auditory thresholds were elevated in mutants, and correlated with cochlear epithelial cell losses. Thus, *Slc26a9*^*P2ACre*^ provides a useful complement to existing pan-otic CRE drivers, particularly for postnatal analyses.

**Summary statement:** We describe a new pan-otic CRE driver, *Slc26a9*^*P2ACre*^, with little activity in the brain or middle ear, and demonstrate its utility by manipulating FGF signaling and assessing hearing loss.

## Introduction

Tg(*Pax2-cre*)^1Akg^, an unmapped BAC transgenic mouse strain expressing CRE recombinase in a pattern that largely recapitulates *Pax2* expression except in optic tissues, is a mainstay of inner ear genetic analyses. Starting at E8.5, it reliably recombines *LoxP* sites throughout the otic placode lineage, which includes the otic epithelium and ganglion neurons, allowing pan-otic conditional deletion or activation of genes from the earliest stages of inner ear development (Ohyama and Groves, 2004). The utility of this strain for postnatal studies of otic genes, however, can be limited when modulating the expression of a given gene-of-interest in an extra-otic portion of the CRE lineage affects viability. For example, we studied embryonic inner ear development following inactivation of *Fgf10* in the Tg(*Pax2-cre*) lineage (Urness et al., 2018), but could not analyze such conditional knockout (CKO) mice postnatally because of P0 lethality, which could be caused by loss of *Fgf10* in the developing kidney (Michos et al., 2010), a component of the Tg(*Pax2-cre*) lineage (Ohyama and Groves, 2004). Similarly, we have been unable to study postnatal *Fgf8*;Tg(*Pax2-cre*) CKO mice due to P0 lethality (unpublished data) suspected to be caused by the requirement for *Fgf8* in the developing mid/hindbrain (Chi et al., 2003), another component of the Tg(*Pax2-cre*) lineage (Ohyama and Groves, 2004) or in the kidney (Perantoni et al., 2005). The targeted *Foxg1*^*Cre*^ allele provides another option for recombining conditional alleles throughout the otic placode lineage (Hebert and McConnell, 2000), but it also has an expansive extra-otic lineage that can prevent postnatal analyses. Thus, *Fgf8*;*Foxg1*^*Cre*^ CKOs also die at birth (Jacques et al., 2007; Zelarayan et al., 2007). Furthermore, *Foxg1*^*Cre*^ is subject to misexpression in some genetic backgrounds (Hebert and McConnell, 2000) and can have genetic interactions as a heterozygote with other otic genes (Pauley et al., 2006). In addition, even if Tg(*Pax2-cre*) or *Foxg1*^*Cre*^ conditional loss- or gain-of-function mice survive postnatally, it can be difficult to unambiguously ascribe auditory or vestibular phenotypes to the inner ear because both *Cre* lineages also include mid- and hindbrain derivatives, including the dorsal cochlear nucleus, inferior colliculus and the cerebellum (Hebert and McConnell, 2000; Ohyama and Groves, 2004; Zuccotti et al., 2012). Furthermore, Tg(*Pax2-cre*) drives CRE expression in pharyngeal arch 1 (Lu et al., 2012; Ohyama and Groves, 2004), which gives rise to middle ear ossicles, and *Foxg1* is expressed in pharyngeal endoderm, which gives rise to the Eustachian tube and tympanic cavity (Anthwal et al., 2013; Chapman, 2011; Sienknecht, 2013). Thus, it is possible that the auditory effects of modulating gene expression using one of these *Cre* drivers might have a middle ear origin.

To expand the options for gene manipulation in the otic placode lineage, we sought to target *Cre* to a gene that is active in the otic placode, but not in the brain, and has extra-otic expression that differs from *Pax2* and *Foxg1*. We noticed that *Slc26a9*, a gene we identified as an otic placode-expressed target of FGFR2b ligands, FGF3 and FGF10, appeared specific to the otic placode as assessed by in situ hybridization of whole E8.5 embryos (Urness et al., 2010). *Slc26a9* is located on chromosome 1 and encodes a membrane protein expressed in adult stomach and lung epithelia that functions as a chloride transporter (Walter et al., 2019). Null mutants are viable and fertile with defects in gastric acid secretion (achlorhydria) and airway clearance in response to inflammation (Anagnostopoulou et al., 2012; Chang et al., 2009; Xu et al., 2005; Xu et al., 2008). We show here that *Slc26a9* is specific for the otic placode, cup and vesicle from E8.5-E10.5. We used *Easi*-CRISPR (Quadros et al., 2017) to insert *P2ACre* into the *Slc26a9* stop codon, producing a correctly targeted allele designed to express CRE from the same transcript as SLC26A9. Reporter analyses of embryonic and early postnatal *Slc26a9*^*P2ACre*^ lineages showed that recombination in otic tissues starts at E9.5, one day later than Tg(*Pax2-cre*), but nevertheless, encompassed the entire otic epithelium and ganglion neurons. There was very little recombination in the brain or middle ear, but perinatal CRE activity was detected as expected in stomach and lung epithelia, as well as in pancreas, kidney and intestinal epithelia. In contrast to *Fgf10* or *Fgf8*;Tg(*Pax2-cre*) CKOs, *Fgf10* or *Fgf8*;*Slc26a9*^*P2ACre*^ CKOs survived postnatally. Auditory brainstem response thresholds were elevated in both CKOs. We also used *Slc26a9*^*P2ACre*^ to drive activation of a dominant-negative FGFR2b ligand trap from E17.5-P3, thereby briefly inhibiting both FGF3 and FGF10 function in the otic lineage, and observed partially penetrant hearing loss correlating with a reduction in the cochlear outer sulcus. These data indicate that *Slc26a9*^*P2ACre*^ mice provide a valuable alternative to Tg(*Pax2-Cre*) and *Foxg1*^*Cre*^, particularly for postnatal analyses of the inner ear, and may also be useful for gene regulation in other FGFR2b-dependent epithelia.

## Results

### *Slc26a9* is otic-specific from E8.5-E10.5, but is not required for auditory function

At E8.5, *Slc26a9* is an otic placode-specific gene dependent upon FGF3 and FGF10 signaling (Urness et al., 2010). To further investigate *Slc26a9* expression we conducted in situ hybridization to whole mount embryos at additional stages (Fig. 1). As the otic placode matured from the 7-8-somite stage (Fig. 1A) through the otic cup (Fig. 1B) and vesicle stages (Figs. 1C-D), *Slc26a9* expression became dorsally regionalized within the vesicle by E10.0-E10.5 (Figs. 1E,E’,F,F’), but remained otic-specific, unlike either *Pax2* or *Foxg1*, which are also expressed in neural and other tissues from early stages (Hebert and McConnell, 2000; Ohyama and Groves, 2004). Notably, we did not detect expression in the otic ganglion. We conducted in situ hybridization to inner ear sections at later stages (E12.5, E14.5, E17.5 and P3), but did not detect a specific *Slc26a9* signal, suggesting that *Slc26a9* otic expression is downregulated after E10.5-11.5. To assess possible requirements for *Slc26a9* in the inner ear, we intercrossed mice heterozygous for an *Slc26a9* null allele (Xu et al., 2008), observed posture/movements (data not shown) and measured auditory brainstem response (ABR) thresholds, an indicator of hearing, to several stimuli at P21-P49 (Fig. 1G). No significant differences between wild type, heterozygous or homozygous mutant mice with respect to behavior or auditory thresholds were detected, showing that *Slc26a9* is not uniquely required during inner ear development or function.

**Figure 1.**
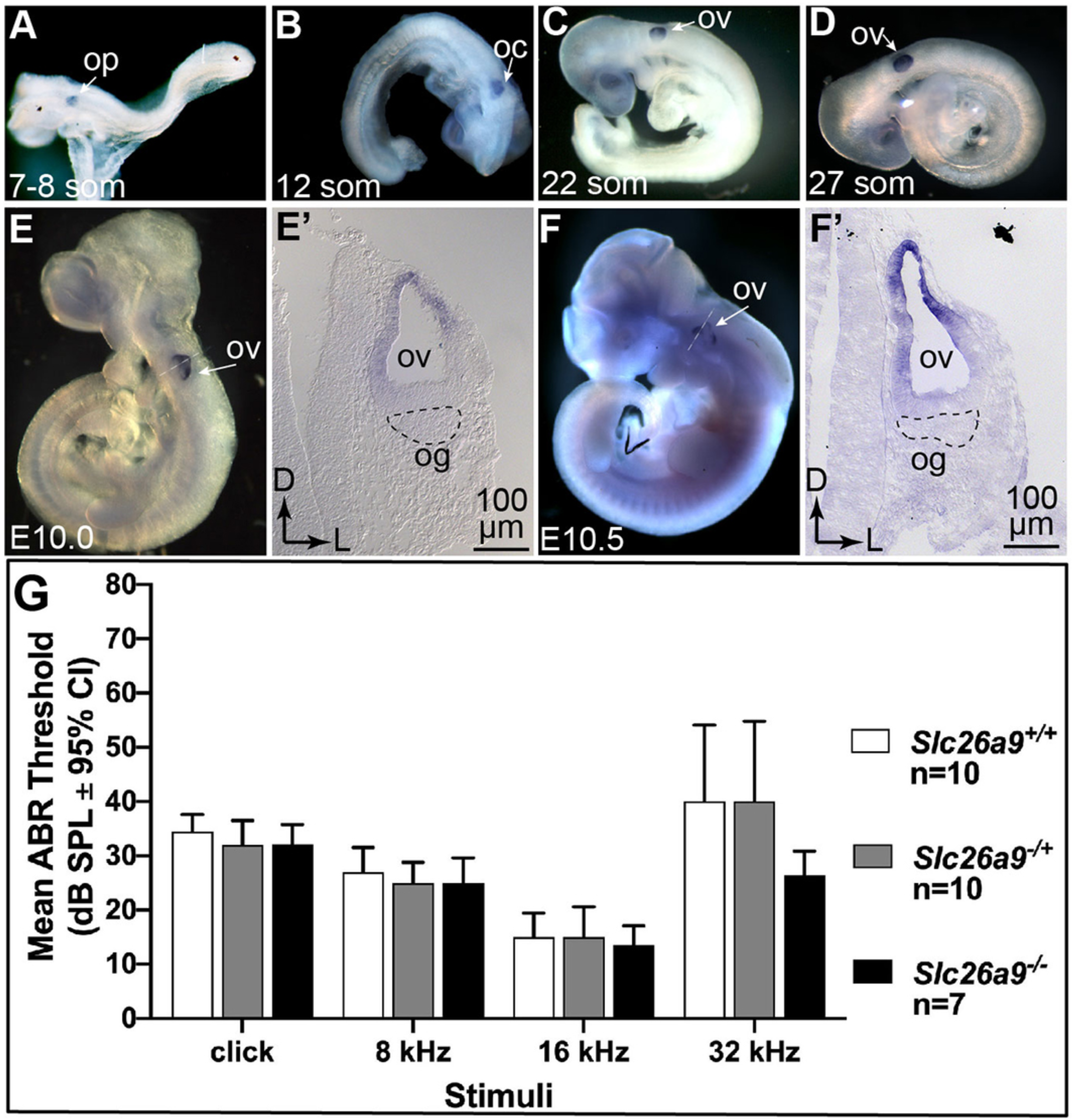
Early expression of *Slc26a9* is otic-specific, but it is not required for auditory function. (A-F) *Slc26a9* mRNA was detected by in situ hybridization to whole mount embryos at the indicated stages. (E’,F’) Transverse sections taken through the otic region of embryos shown in E and F. Dashed lines surround the og. Expression was found throughout the otic placode (A), cup (B) and vesicle (C,D), but was dorsally regionalized by E10.0-E10.5 (E,F). (G) Mean ABR thresholds of P21-P49 *Slc26a9* wildtype, heterozygous and null mice showed no statistically significant differences between genotypes (2-way repeated measures ANOVA with Tukey’s multiple comparisons test). Abbreviations: op, otic placode; oc, otic cup; og, otic ganglion, ov, otic vesicle.

### Targeting *P2ACre* to *Slc26a9*

The substantial differences in extra-otic embryonic *Slc26a9* expression relative to *Pax2* and *Foxg1*, and the lack of a requirement for *Slc26a9* in inner ear development prompted us to target this gene for CRE expression. To avoid disrupting *Slc26a9* expression, we targeted an in-frame *P2A* “self-cleaving peptide” coding sequence (Kim et al., 2011) followed by a codon-optimized *Cre* sequence (Ohtsuka et al., 2013; Shimshek et al., 2002) into the *Slc26a9* stop codon (Fig. 2A-D). In this way, a transcript encoding both SLC26A9 (including 21 extra C-terminal amino acids from P2A) followed by CRE, with an amino terminal proline was expected. We used *Easi*-CRISPR for gene targeting (Miura et al., 2018; Quadros et al., 2017). In this case, a single-stranded DNA donor, together with a ribonucleoprotein complex composed of CAS9, a tracer RNA and a CRISPR RNA designed to cleave inside the *Slc26a9* stop codon, was injected into mouse zygotes. Of 7 founder pups, 4 had the expected 5’ and 3’ junctions with *Slc26a9* (Fig. 2E) Sand these had exactly the expected sequence (data not shown). Founders 6 and 7 were used for germline transmission of the *Slc26a9*^*P2ACre*^ allele and initial crosses of male heterozygotes to female CRE-dependent *LacZ* reporter mice (*Rosa26*^*lslLacZ*^) revealed similar patterns of CRE activity in the offspring, so all further characterization was arbitrarily confined to line 7. When a female heterozygote was crossed to a male *LacZ* reporter mouse, 4 of 5 offspring exhibited ubiquitous ßGAL activity, even though none carried the *Slc26a9*^*P2ACre*^ allele, suggesting that the female’s germline contained CRE, which caused recombination in most zygotes and was propagated to all tissues, similarly to Tg(*Sox2-cre*) (Hayashi et al., 2003). Thus, for all lineage and CKO studies reported here, *Slc26a9*^*P2ACre*^ was contributed by the male parent. Intercrosses of *Slc26a9*^*P2ACre/*+^ mice generated offspring of all three expected genotypes (Fig. 2F). The allele can be maintained as homozygous, with no obvious fertility or longevity decrement for either sex.

**Figure 2.**
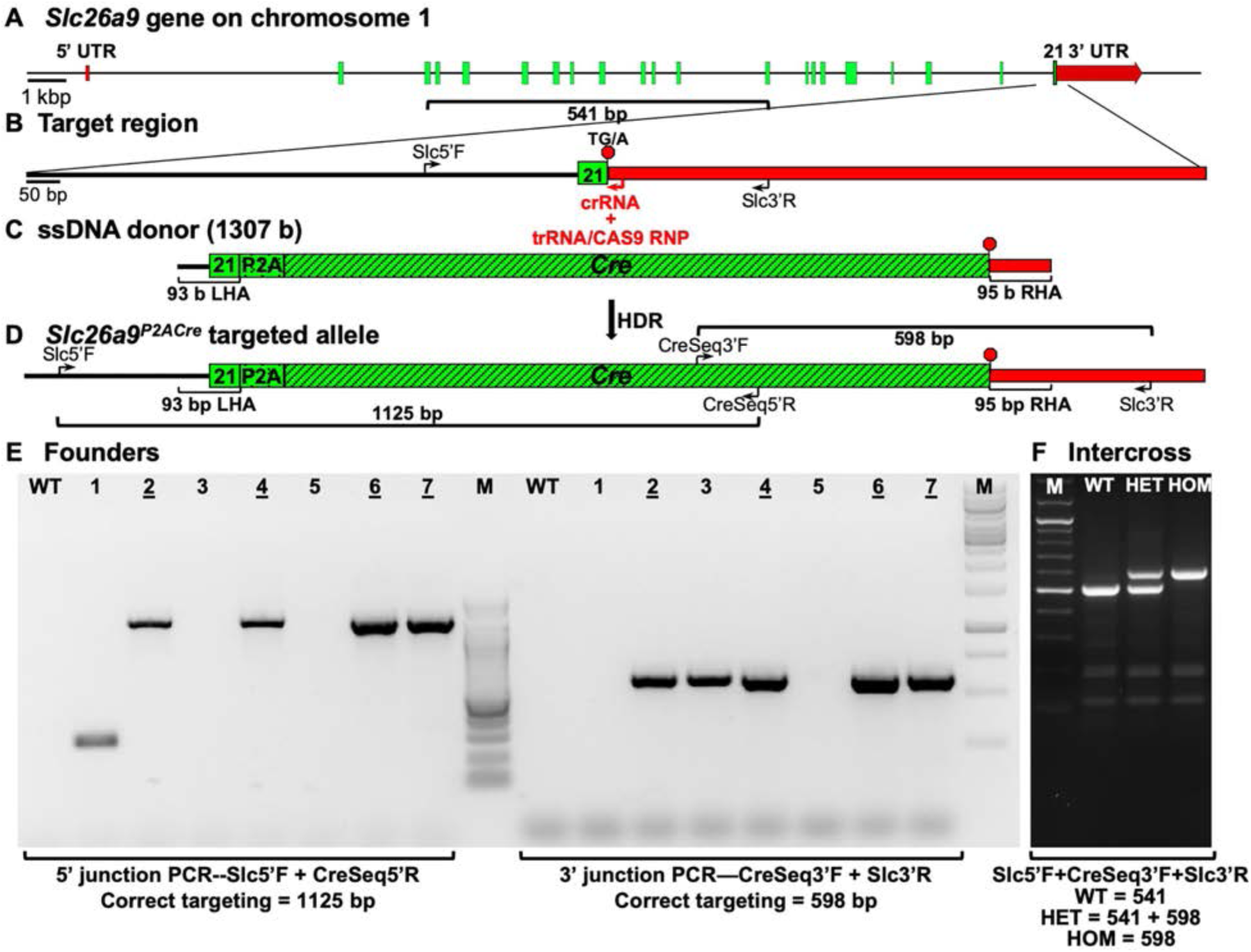
Strategy to target *P2ACre* after the last coding exon of *Slc26a9*, identification of correctly targeted founder mice and implementation of a 3-primer genotyping assay. (A) Schematic of the genomic structure of wild type *Slc26a9*. Exon 21, the last coding exon, is indicated. (B) Enlargement of the 3’ *Slc26a9* region chosen for inserting *P2ACre*. (C) Map of single-stranded DNA (ssDNA) donor sequence used to insert *P2ACre* via *Easi*-CRISPR. (D) Schematic of the 3’ end of *Slc26a9P*^*2ACre*^. (E) Gel separation of PCR products amplified from *Easi*-CRISPR founder pup DNA using 5’ and 3’ primer pairs that encompass the 5’ and 3’ regions of the *Slc26a9* 3’ target region. Sizes of PCR products expected from correct targeting at the 5’ and 3’ junctions of the target site are noted inside brackets below the gels. Identification numbers of pups with correctly targeted PCR product sizes at both 5’ and 3’ ends are underlined (2,4,6,7). DNA sequencing of purified PCR fragments from each of those four founder pups showed the expected sequence (i.e. no mutations). (F) PCR genotyping of 3 pups from an *Slc26a9*^*-/*+^ intercross litter. Expected PCR product sizes for wildtype, heterozygous and mutant DNAs are indicated below the gel. Black lines, green boxes and red boxes indicate non-coding, coding and untranslated mRNA sequences respectively. Black arrows indicate the location and 5’-3’direction of PCR primers. Red arrow indicates the location and direction of the CRISPR RNA (crRNA) used to direct genomic cutting inside the *Slc26a9* stop codon by an RNP complex composed of the crRNA, a universal tracer RNA (trRNA) and CAS9. Abbreviations: LHA, left homology arm; RHA, right homology arm; HDR, homology directed recombination.

### *Slc26a9*^*P2ACre*^ directs CRE activity in the otic lineage starting at E9.5

We surveyed *Slc26a9*-directed CRE activity in sagittal sections of *Slc26a9*^*P2ACre/*+^;*Rosa26*^*lsltdTom/*+^ heads at P1 (Fig. 3). tdTom was found throughout both the cochlear and vestibular epithelia of the inner ear (Figs. 3A,B). Magnified views showed that virtually all otic epithelial cells were labeled, with only a few tdTom-negative gaps (Figs. 3C,C’,D-F). Cochlear (spiral) and vestibular ganglion neurons were also tdTom-positive (Figs. 3A,C,C’,J,K) and their central processes, which form the 8^th^ nerve, were evident (Fig. 3A). Labeling of spiral ganglion neurons was assessed by immunostaining for PROX1. The vast majority of PROX1-positive spiral ganglion neurons were also tdTom-positive (Fig. 3J). Vestibular ganglion neurons were also labeled with tdTom, however, as PROX1 was expressed at much lower levels than in spiral ganglion neurons, we examined co-labeling with antibodies directed against neurofilament (NFH). Virtually all NFH-positive cell bodies in the vestibular ganglion were also TdTom-positive (Fig. 3K). Unlike Tg(*Pax2-cre*), which is active in the derivatives of the mid-hindbrain junction, including the dorsal cochlear nucleus and cerebellum (Ohyama and Groves, 2004; Zuccotti et al., 2012; data not shown), except for the incoming 8^th^ nerve axons, we did not detect the *Slc26a9*^*P2ACre*^ lineage in those structures (Figs. 3A,B), or indeed in any brain nuclei (Fig. 3G and data not shown). There were, however, scattered tdTom-positive cells in the brain that had the appearance of microglia (data not shown). In addition, the anterior pituitary and Eustachian tube were tdTom-positive (Fig. 3G). Finally, although a few tdTom-positive cells were seen in the manubrium of the malleus and in the tympanic membrane (middle ear components, Fig. 3H), these were much less numerous than in sections of P3 Tg(*Pax2-cre*) heads (Fig. 3I).

**Figure 3.**
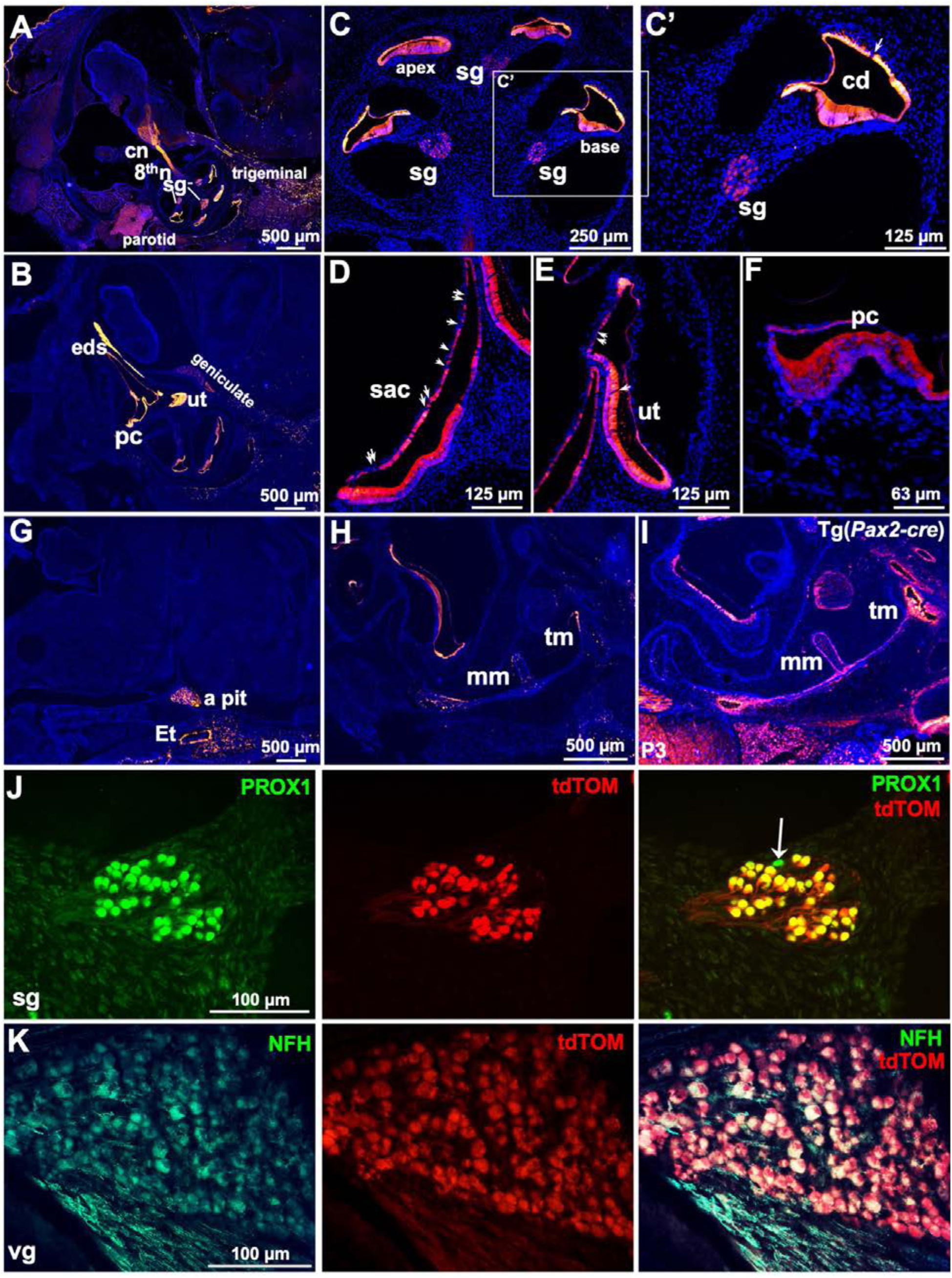
The *Slc26a9*^*P2ACre*^ lineage is found throughout the otic epithelia and ganglia, but is negligible in the brain and middle ear at P1. *Slc26a9*^*P2ACre/*+^;*Rosa26*^*lsltdTom/*+^ P1 heads sectioned in the sagittal plane and counterstained with DAPI. (A,B) tdTom was detected throughout both the cochlear and vestibular epithelia as well as in the cochlear (spiral) ganglion neurons and their processes extending into the cochlear nucleus. Magnified views of the cochlea (C,C’), saccule (D), utricle (E), and posterior canal crista (F) with infrequent tdTom-negative gaps indicated by arrows. (G) The brain was largely devoid of tdTom-positive cells, but the anterior pituitary was labeled. (H) The manubrium of the malleus and tympanic membrane were only sparsely labeled by *Slc26a9*^*P2ACre*^, but were robustly labeled by Tg(*Pax2-cre* (I). (J) Spiral ganglion neurons co-labeled with PROX1 and tdTom, arrow indicates a rare neuron lacking CRE activity. (K) Vestibular ganglion neurons co-labeled with Neurofilament (NHF) and tdTom. Abbreviations: a pit, anterior pituitary; cn, cochlear nucleus; eds, endolymphatic duct and sac; Et, Eustachian tube; mm, manubrium of the malleus; n, nerve; pc, posterior crista; sac, saccule; sg, spiral ganglion; tm, tympanic membrane; ut, utricle; vg, vestibular ganglion.

To determine when *Slc26a9*-directed CRE activity was first detectable, we examined *Slc26a9*^*P2ACre/*+^;*Rosa26*^*lslLacZ/*+^ embryos stained with X-gal. Somewhat surprisingly, there was no ßGAL activity detected at otic placode or cup stages (Figs. 4A,B), but robust, otic-specific activity was present in the otic vesicle and ganglion at E9.5 and E10.5 (Figs. 4C,D). By E11.5, ßGAL activity was detected not only in the otic lineage, but also in the lung and kidney, and diffusely in the lower head and neck (Fig. 4E). Notably, ßGAL activity was not detected in the brain.

**Figure 4.**
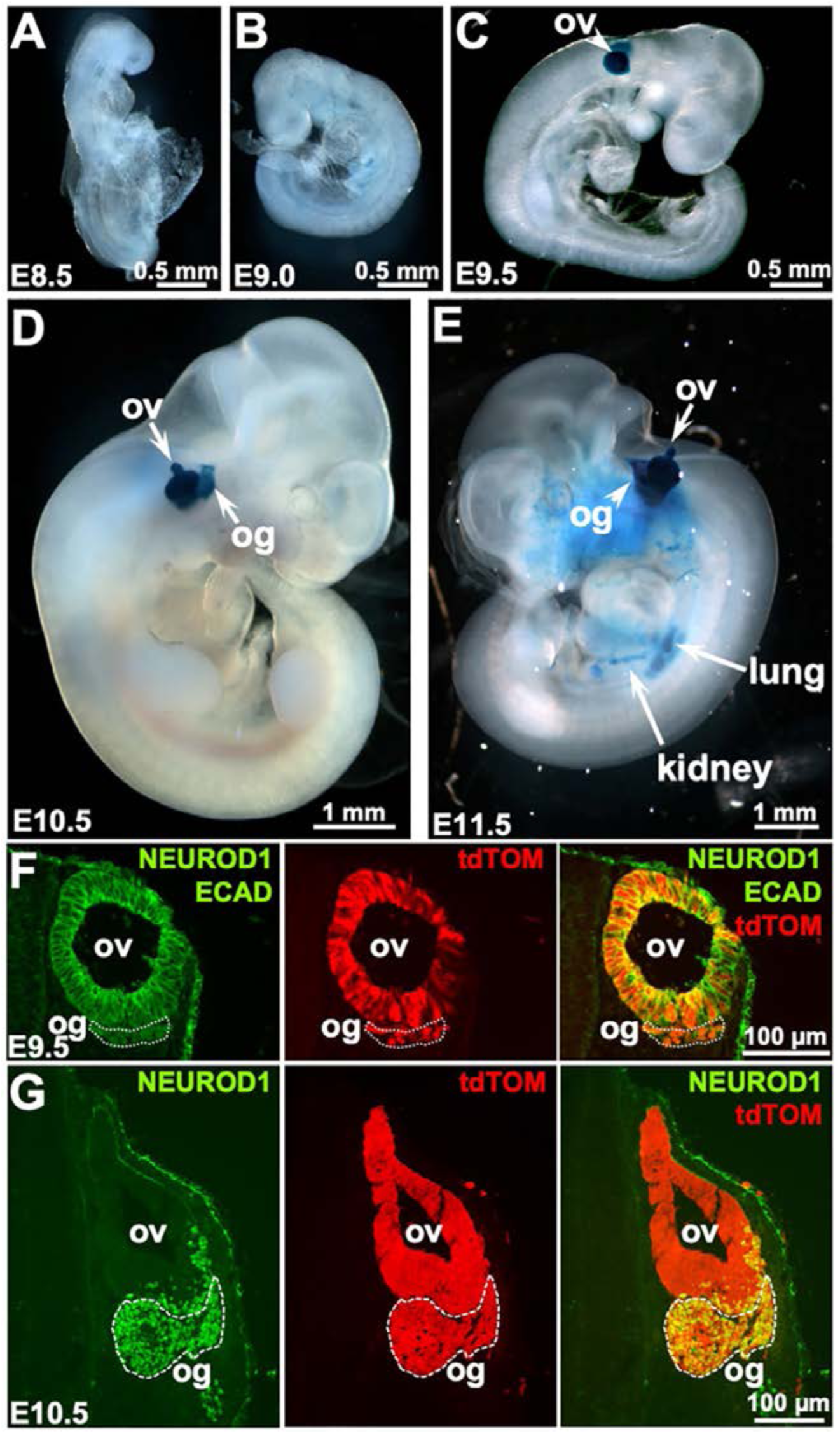
The *Slc26a9*^*P2ACre*^ lineage is first detected at E9.5, and encompasses the otic epithelium and ganglion. (A-E) *Slc26a9*^*P2ACre/*+^;*Rosa26*^*lslLacZ/*+^ embryos stained with X-gal. No ßGAL activity was detected in otic placode (A) or cup-stage (B) embryos. ßGAL was first detected at E9.5 in the otic vesicle (C) and was also evident in the otic ganglion at E10.5 (D). Limited extra-otic expression was apparent by E11.5 in lung and kidney (E). (F,G) *Slc26a9*^*P2ACre/*+^;*Rosa26*^*lsltdTom/*+^ embryos at the indicated stages were sectioned in the transverse plane. E9.5 sections were immunostained with antibodies directed against E-cadherin and NEUROD1 (both in green). tdTom-positive cells (red) were present throughout the otic vesicle and in the adjacent otic ganglion. The merged image shows that neuroblasts (yellow) are present in both the ventral otic epithelium and forming otic ganglion (F). E10.5 sections were immunostained with antibodies directed against NEUROD1. tdTom-positive cells were found throughout the otic vesicle and otic ganglion. The merged image shows double-labeled neuroblasts were still present in the otic epithelium and were abundant in the otic ganglion (G). Ov, otic vesicle; og, otic ganglion.

Since otic lineage labeling was first detected at the otic vesicle stage, which is after the time when otic neurons begin to delaminate from the anteroventral otic cup epithelium (Carney and Silver, 1983; Li et al., 1978), we asked whether all or only a fraction of otic neuroblasts exhibit CRE activity at E9.5 and E10.5. *Slc26a9*^*P2ACre/*+^;*Rosa26*^*lsltdTom/*+^ embryos were sectioned through the otic region and stained simultaneously with antibodies directed against NEUROD1 to detect neuroblasts and E-CADHERIN to outline the otic epithelium. At E9.5 a few NEUROD1-positive cells were detected ventral to the otic epithelium and most appeared to be tdTom-positive (Fig. 4F). By E10.5 much larger numbers of NEUROD1-positive cells had formed a large anteroventral ganglion. The vast majority of these cells were tdTOM-positive (Fig. 4G). Thus, despite the later onset of otic CRE activity in *Slc26a9*^*P2ACre*^ mice relative to Tg(*Pax2-Cre*) mice, cochlear and vestibular ganglion neurons are effectively labeled.

### *Slc26a9*^*P2ACre*^ directs CRE activity in a variety of extra-otic epithelia

To identify additional sites of *Slc26a9*-directed CRE activity we detected tdTom in early postnatal tissues or ßGAL activity in embryos from the reporter crosses. As expected from SLC26A9 expression studies (Amlal et al., 2013; Chang et al., 2009; Lohi et al., 2002; Xu et al., 2005), we detected tdTom-positive cells in sections of early postnatal lung and stomach epithelia, as well as in kidney and pancreas (Fig. 5A-D). It is interesting to note that the gastric lineage includes only the distal (pyloric) portion of the epithelium (Fig. 5B,F). Also, the kidney lineage appears very restricted (Fig. 5C), potentially comprising only the parietal cells of the medullary collecting ducts where SLC26A9 expression has been documented (Amlal et al., 2013). Each of the above mentioned CRE-positive lineages was also detectable in whole mount tissues at E16.5 (Figs. 5E,F,G; pancreas not shown), and the lung epithelium was labeled as early as E11.5 (Fig. 5E, section not shown). Rathke’s pouch, which is the precursor of the anterior pituitary, was strongly labeled at E13.5 and E11.5 (Figs. 5H,L). CRE activity was not assessed in postnatal intestine, but was detected in E13.5 intestinal epithelium (Fig. 5J). Finally, the *Slc26a9*^*P2ACre*^ lineage was quite strong in the E13.5 cornea (Fig. 5K). CRE-negative tissues included heart (Fig. 5E), adrenal gland (Fig. 5G) brain (Fig. 5H), spleen and liver (data not shown).

**Figure 5.**
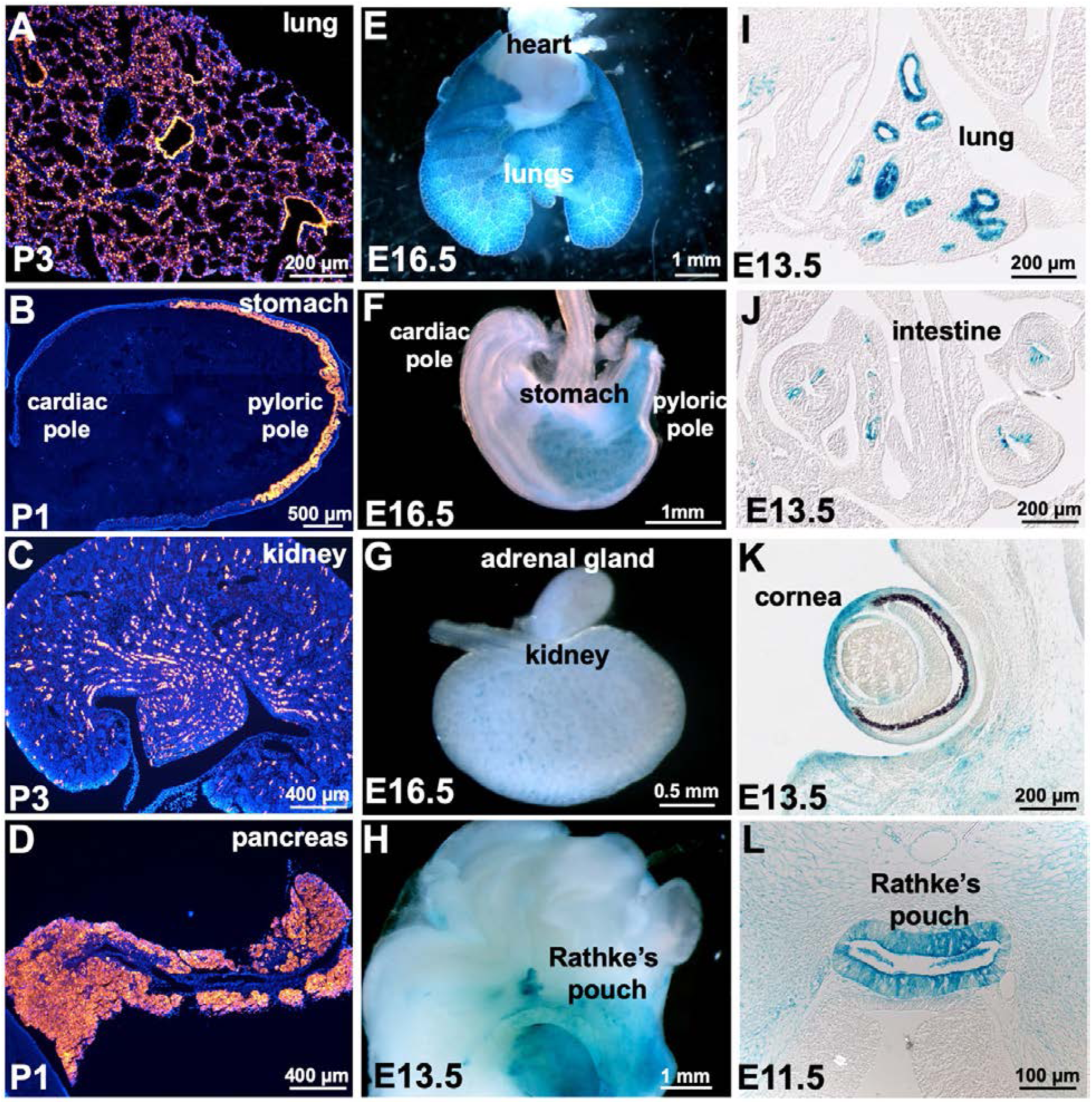
*Slc26a9*^*P2ACre*^-directed CRE activity in selected E13.5-P3 extra-otic tissues. (A-D) *Slc26a9*^*P2ACre/*+^;*Rosa26*^*lsltdTom/*+^ P1 or P3 tissues, sectioned and counterstained with DAPI. (E-H) E13.5 or E16.5 *Slc26a9*^*P2ACre/*+^;*Rosa26*^*lslLacZ/*+^ tissues isolated following X-Gal staining of whole mount embryos. (I-L) E11.5 or E13.5 *Slc26a9*^*P2ACre/*+^;*Rosa26*^*lslLacZ/*+^ tissue sections from X-gal stained embryos.

### *Fgf10*;*Slc26a9*^*P2ACre*^ conditional mutants are viable and show vestibular and auditory dysfunction

*Fgf10* is required for inner ear morphogenesis, but germline null or conditional mutants generated with Tg(*Pax2-cre*) do not survive past birth (Min et al., 1998; Sekine et al., 1999; Urness et al., 2018). To assess postnatal *Fgf10* inner ear phenotypes we generated *Fgf10* conditional mutants using *Slc26a9*^*P2ACre*^ and observed motor behavior and measured ABR thresholds. *Fgf10;Slc26a9*^*P2ACre*^ CKOs survived to weaning in somewhat more than the expected proportion (8/21 = 38%, expected 25%) indicating that the *Slc26a9* lineage does not overlap any sites of *Fgf10* expression required for viability, e.g. the lung mesenchyme (Bellusci et al., 1997; Min et al., 1998) or metanephric mesenchyme (Michos et al., 2010; Ohuchi et al., 2000). 7 of 8 *Fgf10* conditional mutants exhibited mild head tossing or circling behavior and all showed increased ABR thresholds averaging 24-36 decibels above control means for all stimuli. The differences were significant for all comparisons between the CKO and control animals, except at 32 kHz, where variability in two of the control genotypes was high (Fig. 6A).

**Figure 6.**
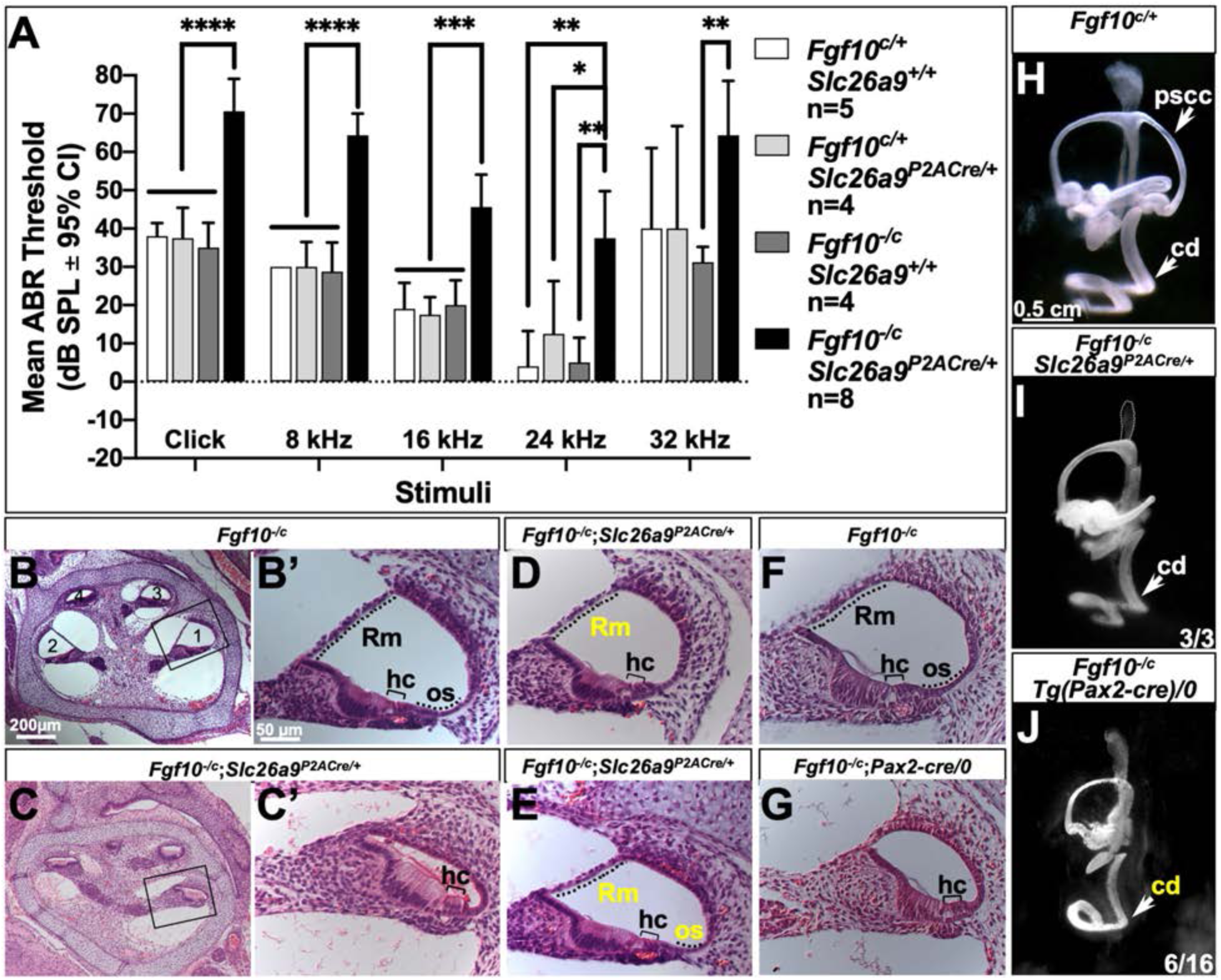
*Fgf10*;*Slc26a9*^*P2ACre*^ conditional mutants have moderate hearing loss and exhibit variable cochlear phenotypes similar to, though milder than, *Fgf10/Tg(Pax2-cre)* CKO mutants. (A) Mean ABR thresholds for *Fgf10*^*c/-*^;*Slc26a9*^*P2Cre/*+^ CKO and *Fgf10*^*c/*+^, *Fgf10*^*c/*+^*;Slc26a9*^*P2ACre/*+^, and *Fgf10*^*c/-*^ control mice at P28-P31. (2-way repeated measures ANOVA, Tukey’s multiple comparisons test; **** adjusted *P*<0.0001, *** adjusted P<0.0002, ** adjusted *P*<0.0021, * adjusted *P*<0.0332). (B-G) H+E-stained sagittal sections through the cochlea of E18.5 *Fgf10*^*-/c*^ (B,B’,F), *Fgf10*^*-/c*^*;Slc26a9*^*P2ACre/*+^ (C,C’,D,E), and *Fgf10*^*-/c*^*;Pax2-Cre/0* samples (G). Boxed basal turns in B and C are shown at higher magnification in B’ and C’. (H-J) Paintfilled ears from control and experimental embryos with genotypes indicated above. The number of ears of each genotype exhibiting the same phenotype is indicated at the lower right. Scale bar in B’ applies to C’-G; scale bar in H applies to I,J. Rm, Reissner’s membrane; hc, hair cells; os, outer sulcus; pscc, posterior semicircular canal; cd, cochlear duct. Yellow text indicates reduction of the structure relative to controls.

To evaluate cochlear development in *Fgf10;Slc26a9*^*P2ACre*^ CKOs relative to previous studies of *Fgf10*^*-/-*^ and *Fgf10;*Tg*(Pax2-cre)* CKOs (Urness et al., 2018; Urness et al., 2015), we examined histologic cross sections of CKO and littermate control cochleae at E18.5 (Fig. 6B-G). The cochlear phenotype of *Fgf10;Slc26a9*^*P2ACre*^ CKOs was variable. One cochlea completely lacked Reissner’s membrane and the outer sulcus (Fig. 6C,C’), and was similar to all of 3 *Fgf10*^*- /-*^ or 4 *Fgf10*;Tg(*Pax2-cre*) CKO cochleae examined previously (Fig. 6G and Urness et al., 2015; Urness et al., 2018). Another cochlea had a shortened Reissner’s membrane and lacked the outer sulcus (Fig. 6D), and a third cochlea had a shortened Reissner’s membrane as well as a reduced outer sulcus (Fig. 6E).

To assess vestibular development in *Fgf10;Slc26a9*^*P2ACre*^ CKOs we paintfilled a few inner ears from the conditional cross and categorized phenotypes according to the scheme used previously for *Fgf10*^*-/-*^ and *Fgf10;*Tg(*Pax2-cre*) CKOs. Relative to controls (Fig. 6H), all 3 *Fgf10;Slc26a9*^*P2ACre*^ CKO inner ears entirely lacked the posterior semicircular canal (pscc; Fig. 6I). This was the most prevalent phenotype observed in *Fgf10;*Tg(*Pax2-cre*) CKOs (8/16; Urness et al. 2018), but in that case we also observed some more severely affected inner ears that lacked the pscc and had an obviously shortened cochlear duct (6/16; Fig. 6J). Thus, histologic and paintfill analyses show that the *Fgf10;Slc26a9*^*P2ACre*^ CKO inner ear structural phenotype is very similar to, but appears milder than that of *Fgf10;*Tg(*Pax2-cre*) CKO inner ears.

### *Fgf8*;*Slc26a9*^*P2ACre*^ conditional mutants are viable and have mild low-frequency hearing loss associated with defective mid-apical inner pillar cell differentiation

*Fgf8*, which is expressed by inner hair cells (IHCs), is required for complete differentiation of the adjacent inner pillar cells (Jacques et al., 2007; Zelarayan et al., 2007). However, the *Fgf8;Foxg1*^*Cre*^ conditional mutants that were studied die at birth, as do *Fgf8*;Tg(*Pax2-cre*) conditional mutants (data not shown), so auditory phenotypes of such mutants cannot be assessed. We generated *Fgf8*;*Slc26a9*^*P2ACre*^ conditional mutants and found that they survived in the numbers expected from the cross (9/35 were CKOs; expected = 8.75). These conditional mutants had modestly, but significantly, elevated ABR thresholds for click and 8 kHz stimuli relative to the *Cre*-negative control genotype tested (Fig. 7A). To assess IPC differentiation, we stained E18.5 whole mount cochleae with antibodies directed against p75^NTR^, and counterstained with phalloidin to reveal IPCs and HCs, respectively (Fig. 7B). As expected from studies of the *Fgf8*;*Foxg1*^*Cre*^ conditional cross (Jacques et al., 2007; Zelarayan et al., 2007), the apical surface of *Fgf8*^*-/c*^ control cochleae showed continuous basal-to-apical expression of p75^NTR^ between the inner and outer HCs (n=3). Similar results were obtained with the other control genotypes, *Fgf8*^*c/*+^ and *Fgf8*^*c/*+^*;Slc26a9*^*P2ACre/*+^ (data not shown). In contrast, *Fgf8*;*Slc26a9*^*P2ACre*^ CKO cochleae (n=4) had gaps in p75^NTR^ staining indicative of incomplete IPC differentiation. The gaps were more numerous in middle and apical segments and had a similar distribution to those seen in *Fgf8*;Tg(*Pax2-cre*)/0 cochleae (n=3). Immunostaining of cochlear cross-sections with antibodies directed against p75^NTR^ and MYO7A (n=2 each) was consistent with these results, showing that in the base of all genotypes, the IPC head extended above the level of the HCs, but in mid and apical regions of both types of *Fgf8* CKOs, the IPC head was level with the HC surface. In addition, the distance between an IHC and the first OHC was narrow (Fig. S1). Thus, the more severe mid-apical disruption in pillar cell differentiation correlates with the modest low frequency hearing loss.

**Figure 7.**
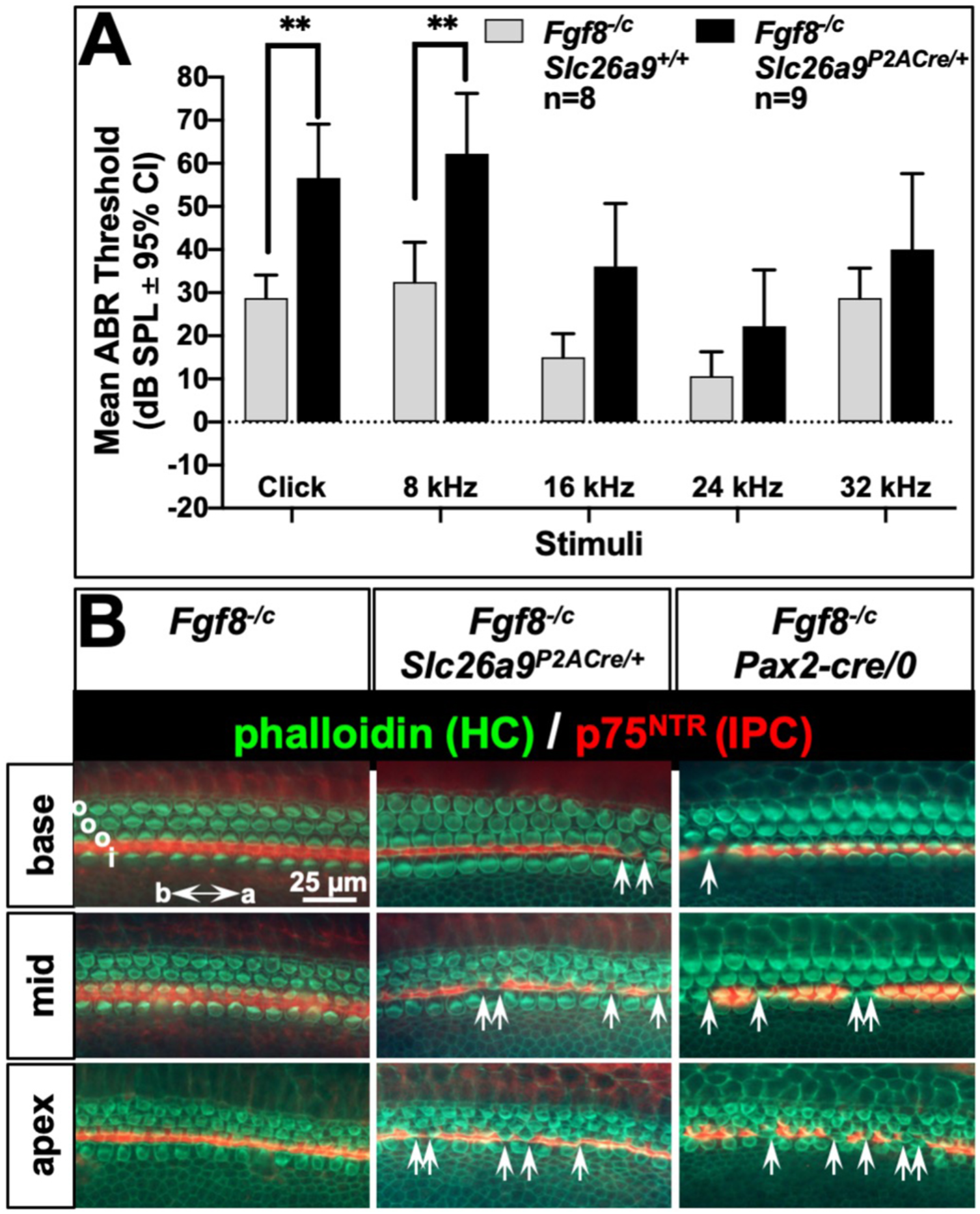
*Fgf8*;*Slc26a9*^*P2ACre*^ conditional mutants have modest low frequency hearing loss and incomplete pillar cell differentiation. (A) Mean ABR thresholds of *Fg8*^*c/-*^;*Slc26a9*^*P2ACre*^ CKO (n = 9) and *Fgf8*^*c/-*^ (n=8) control mice at P31-P50. Error bars show 95% confidence intervals. Asterisks indicate statistical significance of the difference between CKO and control mice (2-way repeated measures ANOVA, Sidak’s multiple comparisons test; ** adjusted *P*=0.0021). (B) E18.5 whole-mount cochleae from control and experimental embryos as indicated above the panels immunostained with phalloidin (green) and anti-p75^NTR^ (red). Apical, mid and basal segments are shown. Arrows indicate gaps in p75NTR immunoreactivity at the epithelial surface, indicative of incomplete inner pillar cell differentiation. i, inner hair cell row; o, outer hair cell row; IPC, inner pillar cell, HC, hair cell.

### Inducing dnFGFR2b from E17.5-P3 in the *Slc26a9* lineage causes hearing loss and disrupts the outer sulcus

To assess the utility of *Slc26a9*^*P2ACre*^ for activating gene expression, we crossed *Rosa26*^*lslrtTA/lslrtTA*^ to *Slc26a9*^*P2ACre/*+^;*Tg(dnFgfr2b)/0* mice and treated pregnant/lactating females from E17.5-P3 with DOX. The cross generated all 4 expected genotypes, of which only one (*Slc26a9*^*P2ACre/*+^;*Rosa26*^*lslrtTA/*+^;*Tg(dnFgfr2b)/0*) was expected to express dnFGFR2b, a ligand trap that inhibits signaling by FGF3 and FGF10 in the inner ear (Urness et al., 2018). Offspring were tested at 3-5 weeks for ABR thresholds. Control pups with any of the 3 genotypes lacking one or both of the components needed to produce dnFGFR2b had normal ABR thresholds for all stimuli, with considerable variability at the highest test frequency (32 kHz; Fig. 8A). In contrast, *Rosa26*^*lslrtTA/*+^*;Slc26a9*^*P2ACre/*+^*;Tg(dnFgfr2b)/0* mice had significantly elevated ABR thresholds (37-44 dB) for all stimuli except 32 kHz, at which the only significant difference was with the corresponding *Cre*-negative control (Fig. 8A).

**Figure 8.**
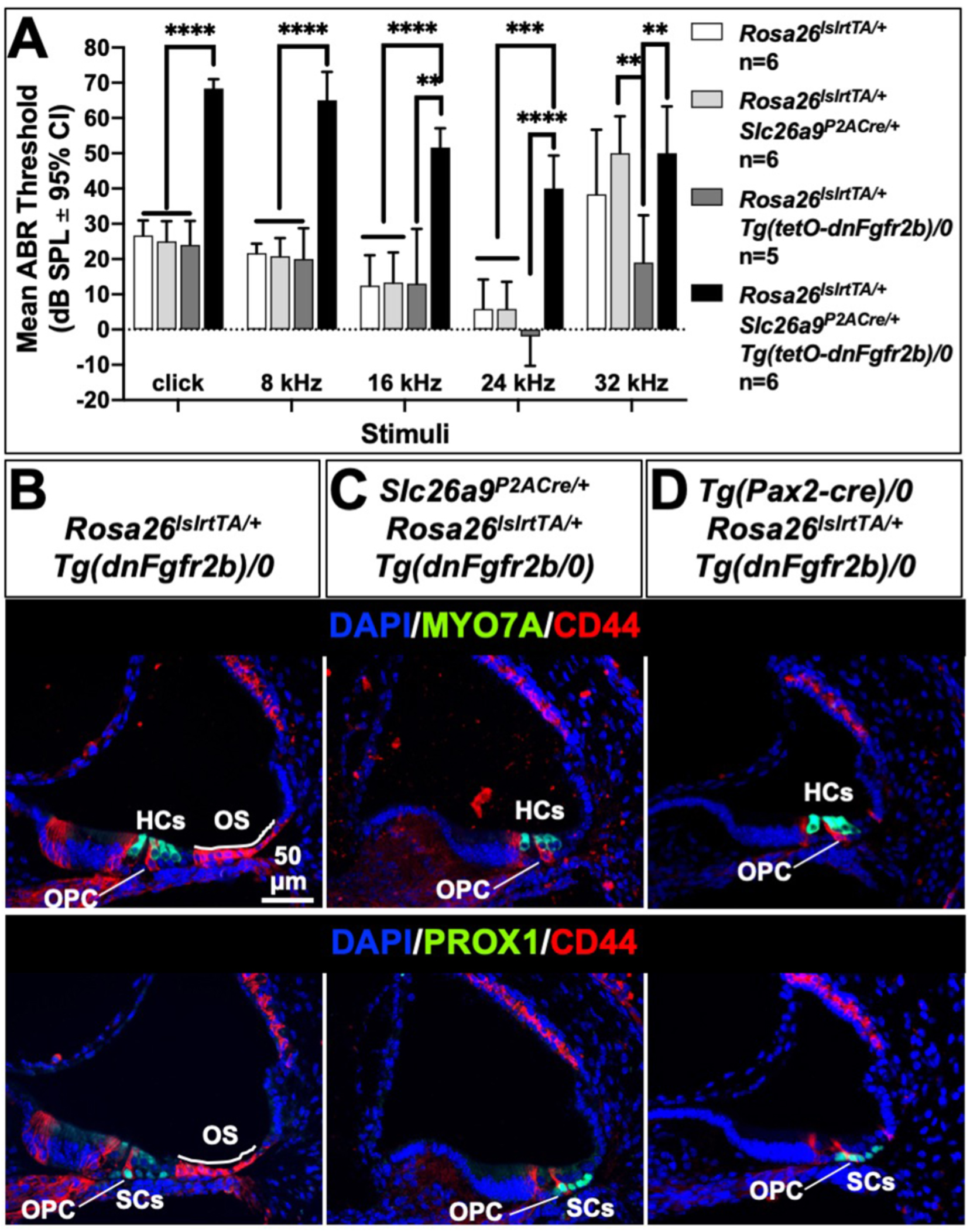
Inducing dnFGFR2b from E17.5-P3 in the *Slc26a9*^*P2ACre*^ lineage causes moderate hearing loss and disrupts the outer sulcus. (A) Mean ABR thresholds of DOX-treated mice of the indicated genotypes at P20-P39. (2-way repeated measures ANOVA, Tukey’s multiple comparisons test; **** adjusted *P*<0.0001, *** adjusted P<0.0002, ** adjusted *P*<0.0021). (B-D) P3 mid-basal cochlear cross-sections of the indicated genotypes stained with MYO7A or PROX1 (green) and CD44 (red) antibodies and counterstained with DAPI (blue). HCs, hair cells; OS, outer sulcus; SCs, supporting cells; OPC, outer pillar cell.

To determine the potential cause of this moderate dnFGFR2b-induced hearing loss, we immunostained P3 cochlear cross-sections with markers of sensory and supporting cells. Normal patterns of MYO7A (sensory hair cells), PROX1 (pillar and Deiters’ supporting cells) and CD44 (outer pillar and non-sensory outer sulcus cells) staining were seen in controls (Fig. 8B; only *Rosa26*^*lslrtTA/*+^;*Tg(dnFgfr2b)/0* is shown, n=2). However, although *Slc26a9*^*P2ACre/*+^;*Rosa26*^*lslrtTA/*+^;*Tg(dnFgfr2b)/0* cochleae had normal sensory and supporting cells, the outer sulcus, a non-sensory tissue required to maintain endolymph composition and that is dependent on *Fgf10* for its specification around E11.5 (Urness et al., 2015) was missing, as indicated by the loss of CD44 staining lateral to the supporting cells (n=3; Fig. 8C). Similar results were seen when *Tg(Pax2-cre)* was used to induce rtTA expression (n=1, Fig. 8D). Thus, *Slc26a9*^*P2ACre*^ readily activates rtTA expression, and FGFR2b ligands are required between E17.5 and P3 to maintain outer sulcus development and normal auditory thresholds.

## Discussion

Our data show that *Slc26a9*^*P2ACre*^ is a promising new tool for otic gene manipulation. Like the classic pan-otic *Cre* alleles, Tg(*Pax2-cre*) and *Foxg1*^*Cre*^, it recombines *LoxP* sites efficiently throughout the otic inner ear epithelia and ganglia, but unlike those *Cre*s, it does not recombine appreciably in the CNS, particularly in the auditory and vestibular nuclei or cerebellum, or in the middle ear ossicles. Thus, it may prove useful for separating peripheral sensory and central origins for auditory and vestibular phenotypes. Furthermore, the extra-otic recombination sites are quite different from those of Tg(*Pax2-cre*) and *Foxg1*^*Cre*^, potentially allowing postnatal survival of conditional mutants, as we found with *Fgf10* and *Fgf8*. This property allowed us to show for the first time that both of these genes are indeed required for normal auditory function.

In comparison with Tg(*Pax2-cre*) CKOs, however, we found that whereas the *Fgf8;Slc26a9*^*P2ACre*^ inner pillar cell phenotypes were similar, the *Fgf10;Slc26a9*^*P2ACre*^ morphologic phenotypes were variably milder than their Tg(*Pax2-cre*) counterparts. Two factors may contribute to this difference. Although *Slc26a9*^*P2ACre*^-directed recombination is pan-otic, it starts at E9.5, one day after Tg(*Pax2-cre*). Since *Fgf8* is not expressed in the cochlea until E15.5 (Hayashi et al., 2007), the deletion with either CRE driver occurs many days before *Fgf8* would normally start to be expressed. Thus, in both conditional mutants, it is unlikely that any *Fgf8* is ever produced in the cochlea and the pillar cell phenotypes are the same. In contrast, *Fgf10* is expressed in a very dynamic pattern starting at or before formation of the first somite in mesenchyme underlying the preplacode and subsequently in the otic cup epithelium starting at E9.0-E9.25 (Pirvola et al., 2000; Urness et al., 2018; Wright and Mansour, 2003). Thus, in both cases, CRE is activated after *Fgf10* has already been produced. The differences between the *Fgf10*;*Slc26a9*^*P2ACre*^ and *Fgf10*;Tg(*Pax2-cre*) CKO phenotypes could reflect a specific requirement for *Fgf10* between E8.5 and E9.5 in subsequent otic morphogenesis and/or perdurance of FGF10 protein after its gene has been inactivated, which is later in the case of *Slc26a9*^*P2ACre*^ than for Tg(*Pax2-cre*), giving more opportunity for FGF10 accumulation. Unfortunately, antibodies able to localize and quantify FGF10 in otic tissues to address this issue more directly have not been described.

Although the basal-to-apical increase in IPC head gaps observed in cochlear surface preparations of *Fgf8*;Tg(*Pax2-cre*) and *Fgf8*;*Slc26a9*^*P2ACre*^ CKOs appeared similar, as did the shortening and narrowing of the IPC space observed in cross-sections, these phenotypes seemed milder than the more frequent and non-graded appearance of IPC head gaps and the cross-sectional IPC area seen in *Fgf8*;*Foxg1*^*Cre*^ CKOs (Jacques et al., 2007; Zelarayan et al., 2007). It is possible that this is a consequence of differing genetic backgrounds. Another possibility, however, is that since *Foxg1* is expressed in the IHC along with *Fgf8*, the previously reported *Fgf8* CKO phenotypes reflect a genetic interaction between *Foxg1*, which is heterozygous in the CKOs, and *Fgf8* to regulate IPC differentiation, similar to the genetic interaction seen between *Foxg1* and *Fgf10* regulating morphogenesis of the inner ear (Pauley et al., 2006). If so, it may be necessary to re-evaluate cochlear CKO phenotypes generated using *Foxg1*^*Cre*^ with this possibility in mind.

One of the characteristics of *Fgf10* germline null cochleae is the absence of both Reissner’s membrane and most or all of the outer sulcus. Based on developmental studies of marker genes in those mutants it appeared that the phenotypes initiated sequentially between E12.5 and E15.5. Thus, it was surprising to find such a severe effect on the outer sulcus after inducing dnFGFR2b starting at E17.5, which is well after the start of outer sulcus specification. This suggests an ongoing requirement for FGFR2b/1b ligands in the maintenance of this tissue; a requirement that is not shared with Reissner’s membrane. The more highly penetrant loss of outer sulcus tissue and somewhat more severe hearing loss following dnFGFR2b induction compared with the *Fgf10* conditional knockout suggests that FGF3, which is expressed just medial to the outer sulcus, presumably in the third row of Deiters’ cells (Kolla et al., 2020; Urness et al., 2018), and which is also a ligand for FGFR2b (Zhang et al., 2006), may be involved in this maintenance function.

The losses/reductions of Reissner’s membrane and the outer sulcus, non-sensory tissues involved in maintaining the unique ionic composition of cochlear endolymph, which is critical for sensory cell function (Jagger and Forge, 2013; Kim and Marcus, 2011; Wangemann, 2006), may explain the reduced ABR thresholds in *Fgf10* CKO and dnFGFR2b-expressing mice. However, the relatively moderate hearing loss found in both cases was somewhat surprising and suggests that the systems for endolymph maintenance are highly redundant. It would be interesting to conduct more detailed studies of the mutant endolymph and determine whether the observed hearing loss progresses in severity over time.

Finally, it is interesting to note that the *Slc26a9*^*P2ACre*^ extra-otic lineages are primarily epithelia that, like the inner ear, are dependent on *Fgf10* for development. For example, *Slc26a9* lineage was found in lung, stomach, intestine, kidney and pancreas epithelia, as well as in Rathke’s pouch, all of which require adjacent *Fgf10* for development (Lv et al., 2019; Min et al., 1998; Ohuchi et al., 2000; Sekine et al., 1999). In addition, corneal epithelial proliferation depends on FGFR2, which is likely stimulated by adjacent mesenchymal FGF10 (Zhang et al., 2015). Thus, not only will *Slc26a9*^*P2ACre*^ be useful for gene regulation in those tissues, but *Slc26a9* expression might serve as a sensor for FGFR2b/1b signaling in many contexts and *Slc26a9* promoter elements could be useful in constructing FGFR2b/1b-dependent reporters.

## Materials and methods

### Mouse alleles, genotyping and crosses

Generation and germline transmission of *Slc26a9*^*P2ACre*^ mice complied with protocols approved by the UNMC Institutional Animal Care and Use Committee (IACUC). Subsequent studies of these and all other mice complied with protocols approved by the University of Utah IACUC. The *Slc26a9*^*P2ACre*^ strain will be submitted to a public repository for distribution. Until then, requests should be directed to Dr. Mansour.

The *Slc26a9* null allele (*Slc26a9*^*-*^; *Slc26a9*^*tm1Sole*^; MGI 3822918) was obtained on a C57Bl/6 background from Dr. Manoocher Soleimani and genotyped using a 3-primer PCR assay modified from Xu et al. (2008). A forward primer in *Neo* (5’TGCGAGGCCAGAGGCCACTTGTGTAGC3’) and a forward primer in exon 5 (5’TGACTGCTGTCATCCAGGTGAGC3’) were combined with a reverse primer in the exon 5-6 intron (5’AAGGGGCATCTGAAAGGTTCTAGGC3’), generating a 325 bp product from the null allele and a 250 bp product from the wildtype allele. Heterozygotes from the first or second generation of backcrossing to CBA/CaJ were intercrossed to generate pups of all three genotypes for ABR testing.

The *Fgf10* null (*Fgf10*^*-*^; *Fgf10*^*tm1.1Sms*^; MGI:3526181) and *Fgf10* conditional (*Fgf10*^*c*^; *Fgf10*^*tm1.2Sms*^; MGI:4456398) alleles and genotyping methods were described previously (Urness et al., 2010; Urness et al., 2015). The *Fgf8* null allele (*Fgf8*^*-*^;*Fgf8*^*tm1Mrc*^; MGI:2150350) was obtained from Dr. Mario Capecchi (Moon and Capecchi, 2000) and genotyped as described (Mansour et al., 2013). The *Fgf8* conditional allele (*Fgf8*^*c*^; *Fgf8*^*tm2Moon*^; MGI:3639320) was obtained from Dr. Anne Moon (Park et al., 2006) and genotyped as described (Fresco et al., 2016). All *Fgf* mutant alleles were kept on a mixed genetic background comprised of C57Bl/6 and various 129 substrains. To generate conditional mutants and controls, *Fgf(8 or 10)*^*c/c*^ females were crossed to *Fgf(8 or 10)*^*-/*+^;*Slc26a9*^*P2ACre/*+^ males.

The origin and genotyping of Tg(*tetO-dnFgfr2b*) (MGI 5582625) was as described (Urness et al., 2018). The *Rosa26*^*lslrtTA*^ allele (Gt(ROSA)26Sor^tm1(rtTA,EGFP)Nagy^; MGI:3583817) was obtained courtesy of Dr. Charles Murtaugh and genotyped as described (Belteki et al., 2005). To enable DOX induction of dnFGFR2b, we crossed *Rosa26*^*lslrtTA/lslrtTA*^*;Tg(tetO-dnR2b)/0* females with *Slc26a9*^*P2ACre/*+^ males. dnFGFR2b expression was induced with an intraperitoneal injection of E17.5 pregnant dams with 0.1 ml/10 g body weight of 0.45 mg/ml (4.5 mg/kg body weight) doxycycline hyclate (Sigma-Aldrich) prepared in PBS, followed by provision of DOX chow (6 g/kg, Custom Animal Diets, LLC) ad libitum until P3, when standard chow replaced DOX chow.

*Tg*(*Pax2-cre*)^1Akg^ (MGI:3046196) substituted for *Slc26a9*^*P2ACre*^ where indicated and was genotyped as described (Urness et al., 2018).

Sex of embryonic and early postnatal samples was not determined. Sex of animals used for ABR measurements was noted and was roughly equal within genotypic classes.

### RNA in situ hybridization

Embryos were harvested and fixed in 4% paraformaldehyde solution (PFA) and stored in methanol at -20°C. Whole mount RNA in situ hybridization of embryos to detect *Slc26a9* mRNA, and fixation, embedding and cryosectioning of stained embryos was conducted as described (Urness et al., 2010). Whole embryos were photographed using a stereomicroscope (Zeiss Discovery V12) fitted with a digital camera (QImaging Micropublisher 5.0). Tissue sections were photographed under DIC illumination (Zeiss Axioskop) using a digital camera (Lumenera Infinity3).

### Auditory Brainstem Response testing

*Slc26a9*^*-/*+^ intercross offspring of both sexes were tested at P21-P49 for auditory brainstem response thresholds to broadband click and 8, 16 and 32 kHz pure tone pip stimuli as described (Mansour et al., 2009). An additional stimulus (24 kHz pure tone pip) was added to the tests of the *Fgf10* (P28-P31) and *Fgf8* (P31-P51) conditional crosses and to DOX-induced dnFGFR2b (P20-P39) offspring. The *Fgf10* CKOs were compared with each of the three control genotypes (*Fgf10*^*c/*+^, *Fgf10*^*c/*+^*;Slc26a9*^*P2ACre/*+^, and *Fgf10*^*c/-*^). The *Fgf8* CKOs were compared with a single control genotype (*Fgf8*^*c/-*^). The *Tg*(*tetO-dnFgfr2b*)/0;*Rosa26*^*lslrtTA/*+^;*Slc26a9*^*P2ACre/*+^ genotype was compared with the *Tg*(*tetO-dnFgfr2b*)/0;*Rosa26*^*lslrtTA/*+^ control genotype. Testing of all pups was performed blind with respect to genotype, but most of the *Fgf10* conditional cross CKO offspring could be identified as such based on head tossing/circling behavior.

Statistical comparisons between genotypes were made using Prism 8.3.1 (GraphPad) to compute 2-way repeated measures ANOVA with Tukey’s or Sidak’s correction for multiple comparisons as indicated to adjust *P*-values.

### Generation of *Slc26a9*^*P2ACre*^ mice

We used *Easi*-CRISPR to generate *Slc26a9*^*P2ACre*^ founders in the C57Bl/6 background as described in detail (Miura et al., 2018; Quadros et al., 2017). Briefly, CAS9 protein, tracer RNA and the *Slc26a9*-specific CRISPR RNA (5’-GGAACCAAAUGGGUCCUCACA-3’) were purchased from Integrated DNA Technologies (IDT) and used to form a ribonucleoprotein complex designed to cleave *Slc26a9* inside the stop codon (TG/A). The long single-stranded donor DNA was custom synthesized by IDT (Megamer) and comprised the 66-base P2A sequence (Kim et al., 2011) in-frame with a codon-optimized *Cre* sequence (Ohtsuka et al., 2013; Shimshek et al., 2002) designed from pBBI (Addgene #6595). These were flanked on the 5’ and 3’ sides with 93 bases and 95 bases, respectively, of homology to *Slc26a9*.

The *Slc26a9*-directed RNP (10 ng/µl) was mixed with single-stranded donor DNA (10 ng/µl) and microinjected into 38 C57Bl/6 zygotes, of which 34 were implanted into the oviducts of two foster mothers following standard procedures described previously (Harms et al., 2014). The seven resulting pups were tested for correct targeting at both 5’ and 3’ sides of the CAS9 cleavage site by PCR amplification of tail DNA using primer pairs Slc5’F plus CreSeq5’R and CreSeq3’F plus Slc3’R. The primer locations and expected PCR product sizes are shown in Figure 2D. The sequences of the primers are: Slc5’F: GGAGGAACACAGTTCACAGT, CreSeq5’R: CTTCTGCCACCTCCCAACTG, CreSeq3’F: GTGTCTGGTGTGGCTGATGACC, Slc3’R: ATGGGTTCACCAGAGTCTCATC. Both PCR products from each of the 4 pups having correctly sized products were sequenced and had no deviations from the expected sequence (data not shown). Two correctly targeted female founders (#6 and #7) were crossed to C57Bl/6 males to transmit the new allele. Male heterozygotes were crossed to *Rosa26*^*lslLacZ/lslLacZ*^ females to assess CRE activity at E10.5, and since both lines gave the same results, only one (#7) was selected for further study. Heterozygous intercrosses were genotyped using a 3-primer mix of Slc5’F+CreSeq3’F+Slc3’R. The wild type and *P2ACre* alleles produce 541 and 598 bp products, respectively. Homozygotes were selected to start a maintenance colony and heterozygotes were used for lineage and conditional crosses.

### Immunostaining of embryonic and postnatal cryosectioned tissues

P1 heads were bisected in the sagittal plane and fixed in 4% PFA, embedded in sucrose/gelatin and cryosectioned at 8 µm as described previously (Mansour et al., 2013). E18.5 or P3 cochleae were isolated and prepared similarly. Whole E9.5 and E10.5 embryos were fixed in 4% PFA, embedded in sucrose/gelatin and cryosectioned in the transverse plane as described (Urness et al., 2018). Primary antibodies were diluted into PBS/5% normal serum of the secondary antibody species/0.2% TritonX-100 and applied at the following dilutions: PROX1 (R&D AF2727, 1:100), NFH (Millipore AB1989, 1:1500), NEUROD (Santa Cruz sc-1084, 1:50), p75^NTR^ (Millipore 07-476, 1:600), MYO7A (Proteus 25-6790, dilution or DSHB 138-1, 1:100), CD44 (BD-Pharmingen 550538, 1:1000). Secondary antibodies (Alexa Fluor^®^ 488 goat anti-rabbit IgG (Invitrogen, A11034), Alexa Fluor^®^ 594 or 488 goat anti-mouse IgG (Invitrogen, A11032 or A11001, respectively) and Alexa Fluor^®^ 594 goat anti-rat IgG (Invitrogen, A11007) were diluted 1:400 into PBST/5% normal serum. DAPI was included in the mounting medium (Vectashield, Vector Labs). Fluorescent signals were observed under epi-illumination on a Zeiss Axioskop and captured using an Infinity3 camera (Lumenera) driven by InfinityAnalyze software. Channels were overlaid using Photoshop CC 2019/2020.

### Histology at E18.5 and paintfilling of E15.5 inner ears

E18.5 heads were bisected in the sagittal plane, fixed in 4% PFA overnight and prepared for paraffin embedding, sectioning and staining with hematoxylin and eosin as described (Mansour et al., 2009). Filling of E15.5 inner ears with latex paint and photography was described previously (Urness et al., 2015).

### Staining of whole mount cochlear surface preparations

Temporal bones were isolated and fixed in 4% PFA overnight and stored in PBS at 4°. Cochleae were freed from the otic capsule and modiolus using fine dissection tools, and Reissner’s membrane, the lateral wall of the cochlear duct, and remnant ganglion were trimmed with fine scissors to reveal the sensory epithelium. Cochleae were rinsed in PBS/0.1% TritonX-100 (PBST), blocked for one hour in 5% normal goat serum in PBST and incubated overnight with rabbit anti-p75^NTR^ (Millipore 07-476, 1:1000) and phalloidin-Alexa Fluor^®^ 488 (Invitrogen A12379, 1:2000). The following day, cochleae were washed three times for 10 min in PBST and incubated with Alexa Fluor^®^ 594 goat anti-rabbit IgG (Invitrogen, A11012) at 1:400 in PBST/5% normal goat serum for 1-2 hours at room temperature. Cochleae were washed as previously described and mounted in 20 µl Fluoromount-G (Invitrogen, 00-4958-02). Fluorescent signals were observed and captured as described above.

## Acknowledgements

We thank Dr. Manoocher Soleimani for providing the *Slc26a9* KO mice, Drs. Scott Rogers, Susan Chapman, Kristen Kwan and Shannon Odelberg, respectively, for consultation on the brain, middle ear, eye and statistics. This work was supported by NIH grants (R01DC011819, R21DC014779, and R01DC014470) to SLM.

## Figure Legends

**Supplemental Figure 1 Inner pillar cell differentiation is affected similarly in *Fgf8*;*Slc26a9***^***P2ACre***^ **and *Fgf8*;*Pax2-Cre* CKOs.** Cochlear cross sections were immunostained to detect hair cells (HC, green) and the inner pillar cell (IPC, red). For all genotypes, basal section IPC heads extend above the level of the HCs. Mid and apical sections of CKOs show IPCs extending only to the level of the HCs, and a narrowing of the distance between the inner hair cell (ihc) and the first outer hair cell (ohc1).

