## Supplemental Figure 1 for "*Slc26a9^P2ACre^*, a new CRE driver to regulate gene expression in the otic placode lineage and other FGFR2b-dependent epithelia"

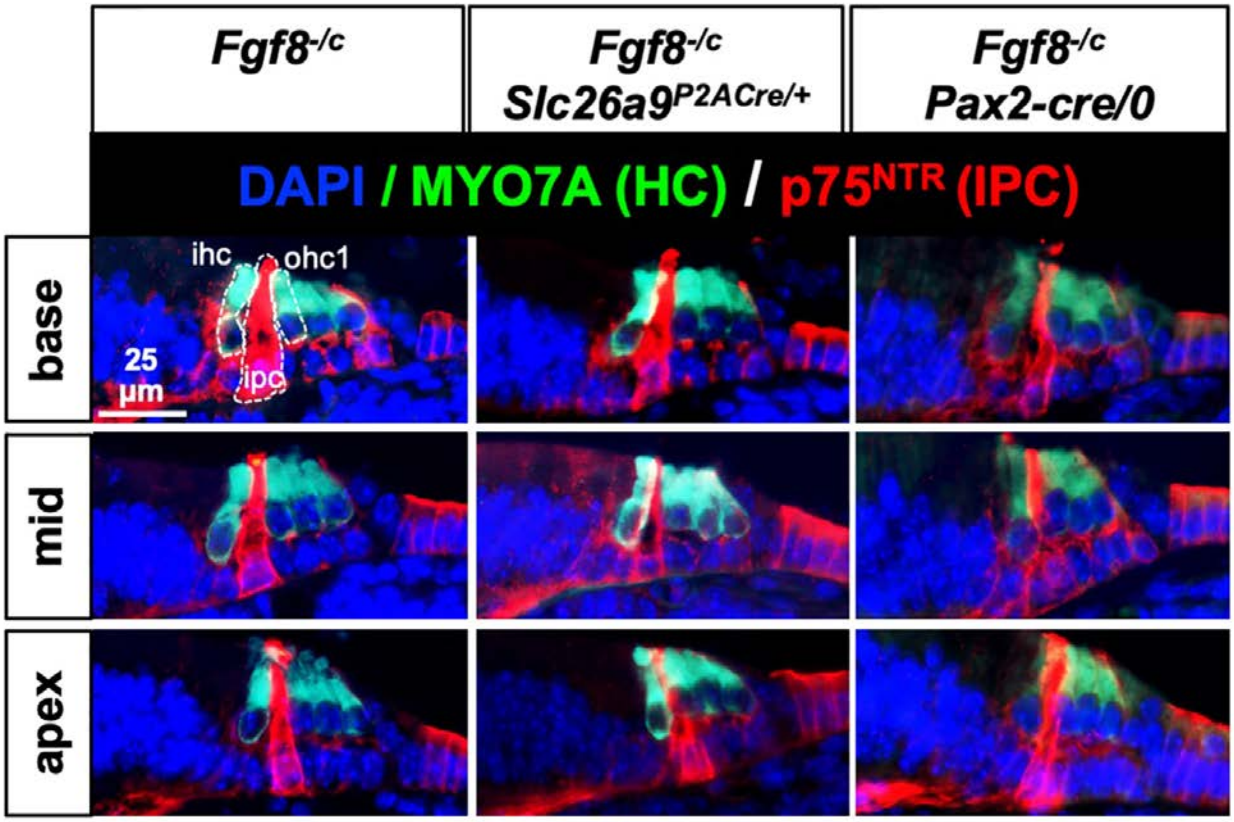


**Supplemental Figure 1 Inner pillar cell differentiation is affected similarly in *Fgf8*;*Slc26a9$^{P2ACre}$* and *Fgf8*;*Pax2-Cre* CKOs.** Cochlear cross sections were immunostained to detect hair cells (HC, green) and the inner pillar cell (IPC, red). For all genotypes, basal section IPC heads extend above the level of the HCs. Mid and apical sections of CKOs show IPCs extending only to the level of the HCs, and a narrowing of the distance between the inner hair cell (ihc) and the first outer hair cell (ohc1).
